# ATG-18 drives longevity in an HLH-30—dependent manner

**DOI:** 10.64898/2026.01.27.701268

**Authors:** Tomas Schmauck-Medina, Alexander Anisimov, David H. Meyer, Yue Hu, Zebo Huang, Yong Wu, Samuel J. Taylor, Michael Takla, Michael R. MacArthur, Sarah J. Mitchell, Ruixue Ai, Anne Simonsen, Vidar Jensen, John Labbadia, Han-Ming Shen, David Rubinsztein, Sofie Lautrup, Guang Lu, Evandro F. Fang

## Abstract

Enhancing autophagy increases lifespan and healthspan in animal models, yet the precise molecular mechanisms underlying these effects are not fully understood. Here we show that overexpression of the essential autophagic gene *atg-18* extends the lifespan of *C. elegans*. We describe a previously unknown, pleiotropic mechanism by which *atg-18* impacts lysosomes and extends lifespan through the transcription factor *hlh-30*, the master regulator of lysosomal biogenesis. We show that, under stress conditions, HLH-30 requires *atg-18* for nuclear translocation. Furthermore, *atg-18* overexpression broadly improves health and stress resilience yet paradoxically increases early-life susceptibility to lethal heat stress. In contrast, enhances heat-stress survival a loss-of-function of function mutation in *atg-18* enhances heat-stress survival, uncovering a temporal-specific effect of *atg-18*. These finding suggest an ATG-18—HLH-30 autophagy—lysosome pathway that plays a key role in lifespan and healthspan.

**Bulleted main discoveries:**

- ATG-18 overexpression increases *C. elegans* lifespan and plays an important role in resilience against starvation and heat stress.
- ATG-18 is required for nuclear translocation of HLH-30
- HLH-30 is required for ATG-18—mediated lifespan extension
- ATG-18 positively regulates lysosomal biogenesis, suggesting a novel role as a molecular coordinator between autophagy and the lysosomal network.

## Introduction

Aging is a complex and multifactorial phenomenon arising from various biological constraints ^1^. Aging research is commonly framed by the hallmarks of aging, a widely used but challenged^2^, description of recurring aging phenotypes. While the original nine hallmarks were proposed in 2013^3^, subsequent research, including our own, has identified additional key features such as impaired macroautophagy^4,5^. Autophagy is an essential cellular process with deep evolutionary roots that orchestrates the degradation of various substrates such as mitochondria, pathogens, and protein aggregates^6,7^. In recent years, the role of autophagy in the process of aging has been highlighted through observations that include age-associated decline of autophagy and its ability to mitigate age-related pathologies through the overexpression of certain autophagic genes in animals, including mammals^8–10^. In addition, the effects on longevity from interventions, such as rapamycin, can be partially blocked when autophagy proteins are downregulated^11^. However, studies have suggested autophagy as a double-edged sword: while autophagy may be beneficial in aging, high levels of autophagy may be detrimental in certain pathologies such as cancer^12^. Furthermore, autophagy may drive age-related pathology in the intestine of *C. elegans*^13^, and neuronal inhibition of some early-acting autophagic genes can increase lifespan^14,15^. Therefore, a crucial task is to identify the conditions under which autophagy is beneficial for health, and those under which it is neutral or harmful.

Here we focus on macroautophagy (hereafter autophagy), a lysosome-dependent pathway in which damaged or superfluous subcellular cargo (e.g., mitochondria) are engulfed by a forming phagophore that elongates and closes to generate a double-membrane autophagosome, which subsequently fuses with a lysosome to enable degradation of the cargo^7^. The classical signature of autophagic membranes is the lipidation of LC3 by conjugation to phosphatidylethalonamine (PE), a process that has been shown to be regulated by the <u>W</u>D-repeat Interacting with <u>P</u>hosphoInositides protein 2 (WIPI2)^16^. In short, during autophagy, the lipid kinase Vps34 assembles into a complex that phosphorylates phosphatidylinositol (PI), producing PI3P^16^. WIPI2 is recruited to the PI3P pools and recruits the Atg12-5-16L1 complex, sequentially driving LC3 lipidation^17^.

In mammals there are four WIPI proteins (WIPI1–4) with shared and distinct functions. WIPI1 and WIPI2 are closely related (∼80% identity), are both induced upon starvation, and colocalize^16^. However, they also exhibit distinct structure localization and non-redundant functional roles; importantly, knockdown of WIPI1 or WIPI2 does not alter expression of the other^16^. Furthermore, WIPI2 exists as multiple isoforms that can differ in recruitment dynamics^17,18^. Curiously, early studies on Atg18 (yeast homologue of WIPI1 and WIPI2) described clusters around the vacuole, the yeast lysosome-equivalent compartment^19^. Later studies found that efficient recruitment of Atg18 to the vacuole requires Vac7 and demonstrated a role for Atg18 in vacuole homeostasis: loss of Atg18 causes enlarged vacuoles and abnormally elevated levels of PI(3,5)P2^20,21^. More recent structural dissections of Atg18 have elucidated the distinct loops at the β-propeller that regulate its recruitment to the forming autophagosome or vacuole^22^.

In mammalian cells, the transcription factor EB (TFEB) is a master regulator of lysosomal biogenesis. During starvation or lysosomal stress, lysosomal Ca^2+^ release activates the phosphatase calcineurin. Calcineurin then dephosphorylates TFEB, enabling its translocation to the nucleus^23^. In the nucleus, TFEB directly binds to the CLEAR motif (a 10-base E-box-like palindromic sequence named Coordinated Lysosomal Expression and Regulation) in the promoter regions of target genes, transcriptionally upregulating a broad set of genes involved in autophagy, lysosomal biogenesis, exocytosis, and endocytosis^23–26^. In contrast, under nutrient-rich conditions, TFEB is phosphorylated at Ser211 by mTORC1, which leads to its binding to 14-3-3 proteins and its effective sequestration in the cytoplasm^27,28^. Moreover, mTORC1-mediated TFEB phosphorylation at Ser142 and Ser138 is critical to control TFEB nuclear export^29^.

Interestingly, *in vivo* studies in *C. elegans* have implicated the TFEB orthologue HLH-30 as a key regulator of longevity. For instance, *hlh-30* is required for the full lifespan extension of several long-lived mutants, including *daf-2, eat-2*, and *glp-1*, among others^30^. In contrast, inhibiting HLH-30 nuclear export induces autophagy and extends lifespan^31^. Similarly, *atg-18* knockdown or loss-of-function drastically shortens wild-type lifespan and diminishes the lifespan extension observed in the long-lived strains mentioned above^32–36^. In mammalian cells, overexpression of WIPI2B has been shown to rescue the neuronal age-related decline in autophagosomal biogenesis^37^. Whether ATG-18 can increase lifespan on its own, what its downstream molecular mechanisms are and its pleiotropic effects, remains unknown. In this study, we describe the outcomes of overexpressing the autophagic gene *atg-18*. We investigated the roles of *atg-18* in aging and fitness using the nematode *C. elegans*, further validating the degree of mechanistic conservation in human cells. Our data suggest that *atg-18* lifespan extension is *hlh-30* dependant.

## Results

### *atg-18* overexpression extends *C. elegans* lifespan and is essential for health

To study whether *atg-18* levels have an impact on survival and health in both physiological and stress conditions, we generated multiple *C. elegans* strains varying in level and spatial-expression of *atg-18*. First, we investigated the effects of both ubiquitous (*atg-18* OE^peft-3^) and neuronal (*atg-18* OE^prab-3^) overexpression of *atg-18* under *eft-3* and *rab-3* promoters, respectively. We found that both ubiquitous and neuronal overexpression of *atg-18* increased the nematodes median lifespan by 21.4% (**Fig. 1a**) and 42.9% (**Fig. 1b**), respectively. Given the pronounced effect of neuron-specific *atg-18* overexpression, we asked whether neuronal rescue alone could restore lifespan in the loss-of-function mutant *atg-18(gk378)*. As expected^38,39^, the mutation of *atg-18* resulted in a 28.6% reduction in median lifespan, and although neuronal rescue significantly extended lifespan, it remained with a median lifespan 21.4% shorter (**Fig. 1c**).

**Fig. 1.**
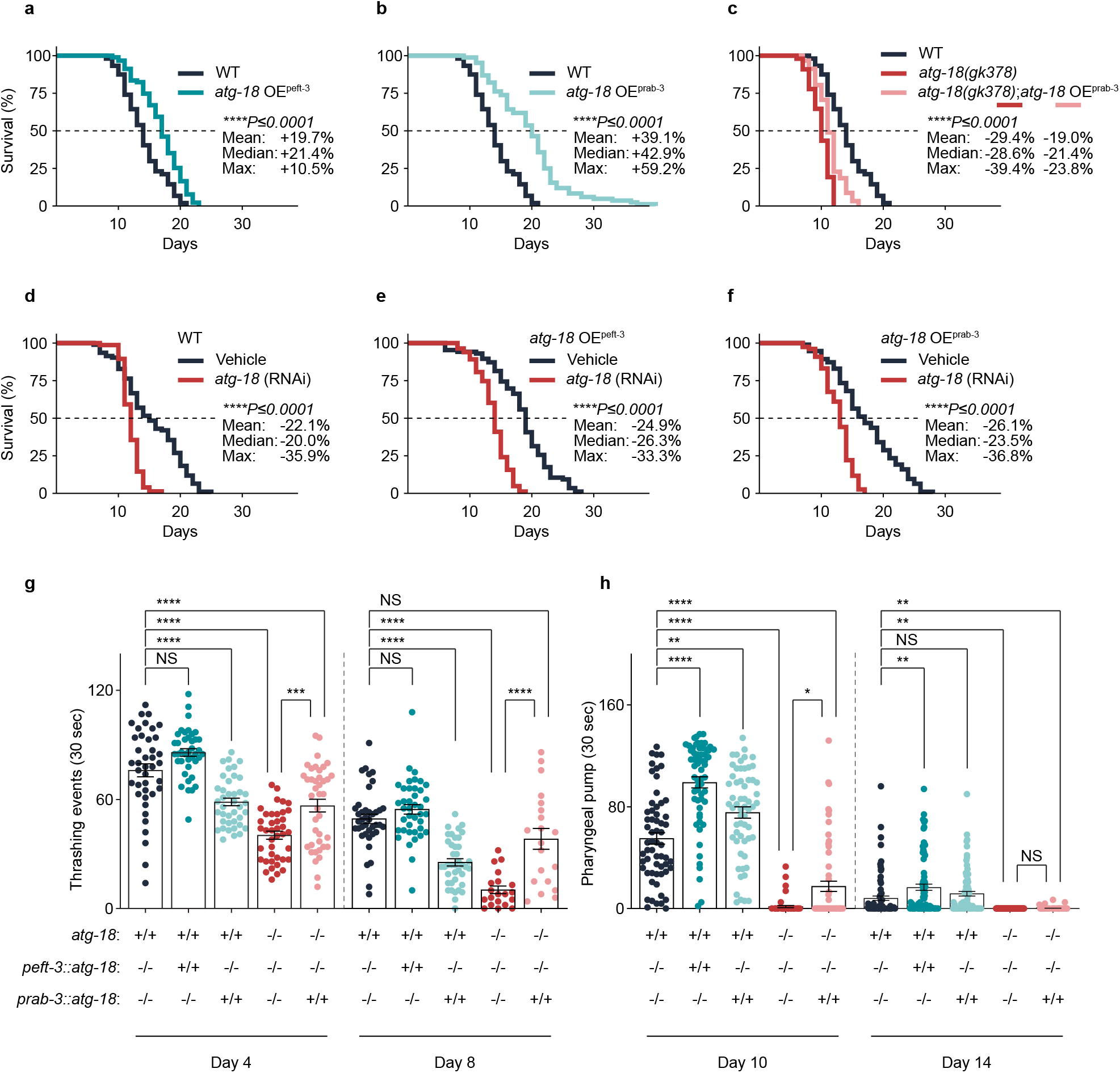
Overexpression of *atg-18 e*xtends lifespan and increases healthspan in *C. elegans*. **a-c**, Lifespan analysis of *atg-18* OE^peft-3^ **(a)**, *atg-18* OE^prab-3^ **(b)** and *atg-18* OE^prab-3^ in the mutant background **(c)**. The effect of *atg-18* RNAi on WT **(d)**, *atg-18* OE^peft-3^ **(e)** and *atg-18* OE^prab-3^ **(f)**. L4440 (empty vector) was used as the vehicle condition. Data represent four biological replicates. ∼100 worms were used for each group/biological repeat. All quantitative data were shown in median ± S.E.M. Log-rank test was used for the statistics of lifespan data. **g**, Swimming score at day 4 and 8 of adulthood **h**, Pumping rate at day 10 and 14 of adulthood. Each graph represents data pooled from three biological replicates. At least 20 animals were scored in each strain at each age.

To confirm that the increased lifespan is driven by *atg-18* rather than off-target effects, we knocked down *atg-18* via RNAi feeding from birth. First, we found that *atg-18* knockdown in WT (wild-type) animals resulted in a 20% reduction of lifespan (**Fig. 1d**). Furthermore, we found that, *atg-18* knockdown reduced the lifespan of *atg-18* OE^peft-3^ animals by 26.3%, essentially abolishing the increased lifespan (**Fig. 1e**). Curiously, the lifespan extension of *atg-18* OE^prab-3^ was less pronounced under control L4440 bacteria than previously observed under OP50 (**Fig. 1b, f**). Unexpectedly, RNAi knockdown of *atg-18* largely abolished the lifespan extension observed in the *atg-18* OE^prab-3^ strain, effectively restoring lifespan to near wild-type levels (**Fig. 1f**). Given the well-documented inefficiency of RNAi in neurons^40^, we did not expect standard RNAi treatment to negate the lifespan extension of the neuronal overexpression strain. However, previous studies have reported that the rab-3 promoter can exhibit leakage outside neuronal tissues, which can be reversed by RNAi^33,41^. Together, these observations suggest that the lifespan extension is unlikely to be driven primarily by increased neuronal *atg-18* expression, but rather by broad elevated *atg-18* expression.

We next asked whether *atg-18* overexpression could improve healthspan by checking a series of endpoints, such as reproductive aging (fecundity and the formation of uterine teratoma-like tumours in germline), body bending (thrashing/swimming), and pumping. Reproductive age is one of the most essential features of evolutionary fitness. Curiously, *atg-18* overexpression did affect fecundity, meanwhile *atg-18(gk378)* mutans exhibited a dramatic reduction in fecundity (**Supp. Fig. 1a**). Furthermore, the neuronal rescue of *atg-18* could not fully rescue the number of eggs laid by *atg-18(gk378)* animals (**Supp. Fig. 1a**). Uterine teratoma-like tumours in the germline of *C. elegans*, bearing similarity to mammalian ovarian teratomas, have been suggested to be driven by quasi-programs of aging^42^. To further investigate the reproductive health of these worms, we looked for the presence these tumours. While at adult Day 1 (D1) there were no significant differences among the strains tested, at D4 all the gonads of *atg-18(gk378)* mutants had ‘teratoma’-like structures, in comparison to only 15% in WT animals (**Supp. Fig. 1b**). In contrast, strains overexpressing *atg-18* did not differ in teratoma formation from WT. Curiously, neuronal rescue of *atg-18* was sufficient to reduce the incidence of teratoma-like structures in *atg-18(gk378)* animals from 100% to 30% (**Supp. Fig. 1b**).

Next, we decided to measure the physical fitness of the animals. The rate of pharyngeal pumping and body movement are two measures that decline with age in *C. elegans*^43,44^. Compared to WT worms, *atg-18* OE^peft-3^ increased swimming rate by 12.8% and 10.6% at Day 4 (D4) and Day 8 (D8) respectively (**Fig. 1g**). They also performed better than WT at pumping rate having 79.6% and 105.6% increase at Day 10 (D10) and Day 14 (D14) respectively (**Fig. 1h**). However, compared with WT animals, *atg-18* OE^prab-3^ reduced swimming performance by 22.9% and 48.4% at D4 and D8 respectively. Finally, neuronal rescue of *atg-18* improved each metric except pumping rate at D14, compared to the loss-of-function *atg-18(gk378)* (**Fig. 1g, h**), which could be explained by the fact that at D14 the differences between these two groups becomes smaller, as most animals have ceased to pump. Overall, these data suggest that *atg-18* plays an important role in physical fitness.

### Trade-off: differential roles of *atg-18* in thermotolerance between early-life and middle-life

In biology, a trade-off refers to an event where an organism improves a biological system at the cost of another. For example, the disposable soma theory of aging proposes that organisms are faced with a trade-off between investing in either development, reproduction or maintenance^45^. This provides an explanation for observations in which long-lived *daf-2* mutants, when co-cultured with WT worms, are driven to extinction over multiple generations as a result of reduced reproductive fitness^46^. Laboratory conditions are quite different to the challenging conditions animals encounter in nature. Therefore evolution has developed key mechanisms to deal with them for survival. Hence, we decided to test multiple stressors on the animals to look for potential trade-offs.

Starvation is one of the most ubiquitous types of stressors in nature. Indeed, mechanisms to deal with energy restriction can be found from yeast to humans. For example, autophagy is a critical mechanism that facilitates survival under scarcity of nutrients^7^. With that considered, we carried out a starvation assay at larval stage L1^47^. We found that *atg-18(gk378)* animals exhibit high vulnerability to starvation (88.3% reduction in survival), while neuronal rescue of *atg-18* dramatically improved the survival of *atg-18(gk378)* animals (56.6% higher survival) (**Supp. Fig. 2a**). We found no significant effect from overexpression of *atg-18* on survival. These data suggest that *atg-18* is essential for survival under starvation.

Another highly conserved biological system is the response to temperature-related shocks. We assessed thermoresilience of worms with different *atg-18* levels by exposing young adults to either cold shock (4°C for 24 hours) or heat shock (37°C for 3 hours), followed by a 24-hour recovery period under 20°C before analysis. Unexpectedly, we found that *atg-18(gk378)* mutants were substantially more resistant to both stress conditions compared to WT animals at D1, exhibiting a absolute 60% higher survival rate following cold shock and a 88% higher survival rate following heat shock (**Supp. Fig. 2b**). Given the possibility that the mutants might be developmentally delayed, thereby introducing a potential confounder, we repeated the heat-shock assay at day 2 of adulthood (D2). Although a trend toward increased resistance in mutants persisted, the difference was no longer statistically significant, attributable to a decline in mutant thermotolerance while WT resistance remained comparable to that observed at D1 (**Supp. Fig. 2b**). Consistent with a potentially negative role for *atg-18* in early-life thermotolerance, both ubiquitous and neuronal *atg-18* overexpression strains exhibited reduced absolute survival at D2 compared to wild type (−14% and −32%, respectively; **Supp. Fig. 2c**). Furthermore, in agreement with these findings, neuronal rescue of *atg-18* diminished stress resistance at D1 of *atg-18(gk378)*, resulting in a 32% reduction in survival following cold shock and a 27% reduction following heat shock (**Supp. Fig. 2b**). Later in life, at day 8 (D8), the thermoresilience of *atg-18(gk378)* animals was dramatically lost (**Supp. Fig. 2d**), with *atg-18(gk378)* mutants having a 35% lower survival compared to WT. By contrast, at this age, both *atg-18* OE^peft-3^ and *atg-18* OE^prab-3^ had superior thermoresilience than WT animals, with survival rates 28% and 35% higher respectively. Furthermore, the neuronal rescue in *atg-18(gk378);atg-18* OE^prab-3^ animals also resulted in increased survival compared to the *atg-18(gk378)* mutants (**Supp. Fig. 2d**).

### ATG-18 has profound effects on transcriptomic profile

To further understand the molecular mechanisms of how *atg-18* impacts aging, we performed RNA sequencing (RNAseq) on all the strains at early-(D1) and middle (D8)-adulthood. There was a great extent of variation in the D1 data (**Fig. 2a**); while the D8 data showed a more consistent pattern (**Fig. 2b**). PCA of D8 revealed different groups, with a clear difference between WT and *atg-18(gk378)*. The neuronal rescue *atg-18(gk378)*;*atg-18* OE^prab-3^ strain exhibits a pattern more similar to WT, suggesting a universal transcriptional benefit by the neuronal expression of *atg-18* (**Fig. 2b**). A hierarchical clustering heatmap analysis based on a correlation metric showed that the chronological age has the strongest impact on the transcriptional status (**Fig. 2c**). Within each timepoint, the five strains were further clustered. Notably, at both timepoints the ubiquitous and neuronal overexpression strain cluster closest together. *atg-18(gk378)* mutants formed a separate cluster at D8, but clusters closest to the neuronal rescue at day 8. Note that at D8, *atg-18(gk378)* mutants, had the most extreme values, indicating the biggest differences. While still clustering with *atg-18(gk378)*, the neuronal rescue shifted the transcriptional profile closer to that of the other groups (**Fig. 2c**).

**Fig. 2.**
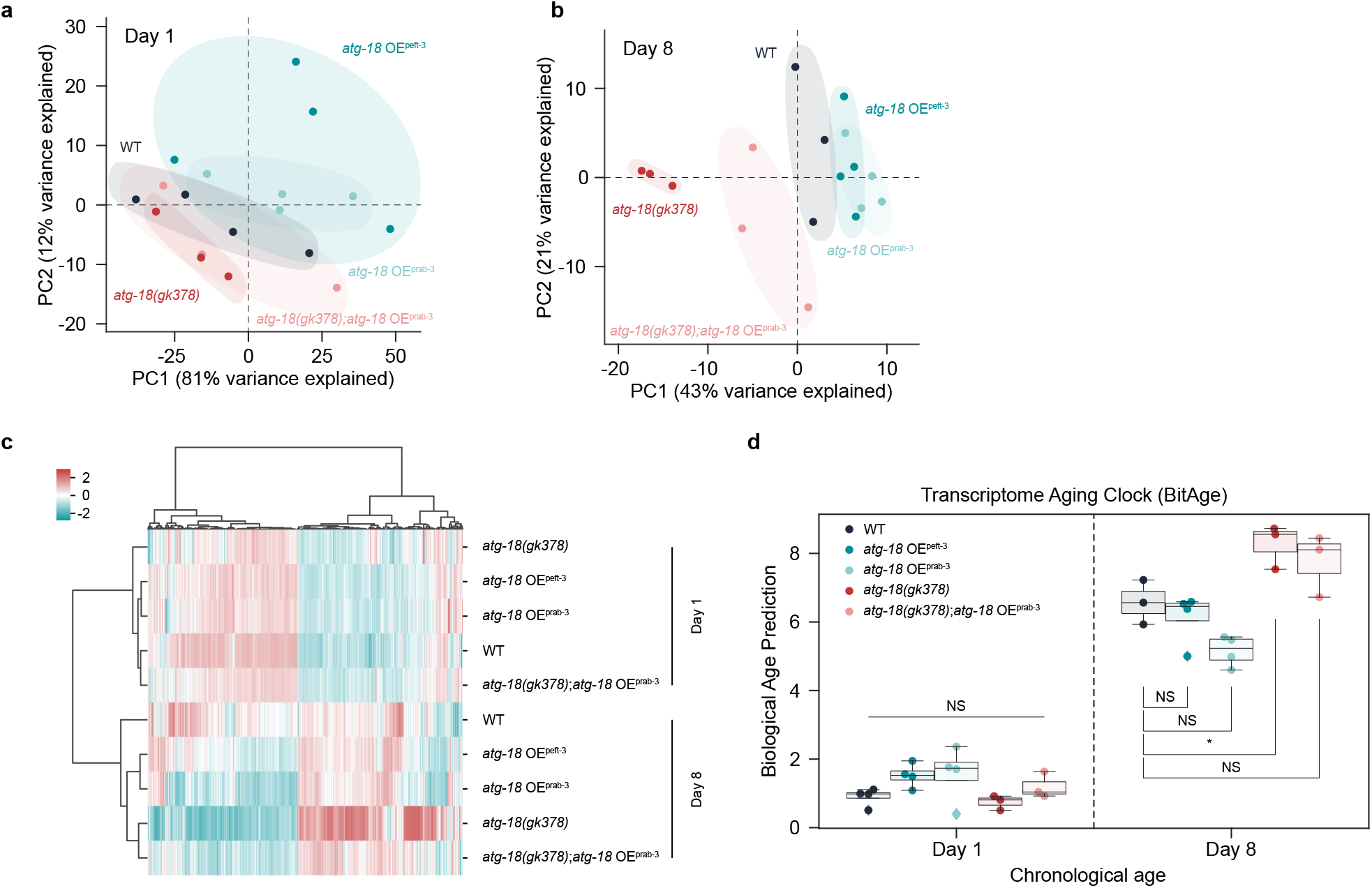
RNA-seq analysis uncovers numerous transcriptional changes. **a-b**, Principal component analysis (PCA) plot of different *atg-18* strains at day 1 and day 8 based on the log transformed gene count tables from the RNAseq analysis. **c**, Heatmap of RNAseq analysis showing gene expression profiles. Each column represents a differentially expressed gene, and each row represents a strain at different ages (D1 and D8). The color gradient reflects the level of gene expression, with red indicating higher expression and blue indicating lower expression. **d**, Biological age (BitAge) prediction based on transcriptomic profiles for strains with different expression levels of *atg-18*. Each strain at every age was analyzed with a sample size of at least 300 animals.

While chronological age does not always reflect ‘biological age’, we asked whether overexpression of *atg-18* could reduce measures of biological age. We applied ‘BitAge’, which is based on transcriptomic profiles and directly predicts not chronological but the biological age of groups^48^. As expected, the differences were mild among the groups in D1 worms while distinct patterns were seen in older age. At D8, *atg-18(gk378)* mutants showed a significantly older biological age compared with WT (**Fig. 2d**); both ubiquitous and neuronal overexpression of *atg-18* showed trends towards a reduced biological age in WT animals (**Fig. 2d**); furthermore, neuronal rescue of *atg-18* trended to a reduction of biological age in the *atg-18(gk378)* mutants (**Fig. 2d**). We suspect that the ages analysed were too early in life to capture large age-dependent effects of *atg-18* expression. Moreover, aging is a complex, multidimensional process that may not be fully captured by current aging clocks or by a single omics datatype. Indeed, similar observations have been reported by others: for example, late *daf-2* knockdown did not result in significant changes in BitAge predictions, indicating that not all biologically relevant age-related patterns are necessarily reflected at the transcriptomic level^49^.

### ATG-18–HLH-30 interdependence in lifespan extension and nuclear translocation

Based on the RNA-seq data, we asked whether *atg-18* affects activities of any transcription factors. Using motif enrichment analysis, we found that in the strains overexpressing *atg-18*, genes with an HLH-30 motif at the promoter region were strikingly upregulated early in adulthood (**Fig. 3a**). On the other hand, *atg-18(gk378)* mutant displayed a significant downregulation of such genes, suggesting a potential role for *hlh-30* as a downstream executor of *atg-18* dependent phenotypes.

**Fig. 3.**
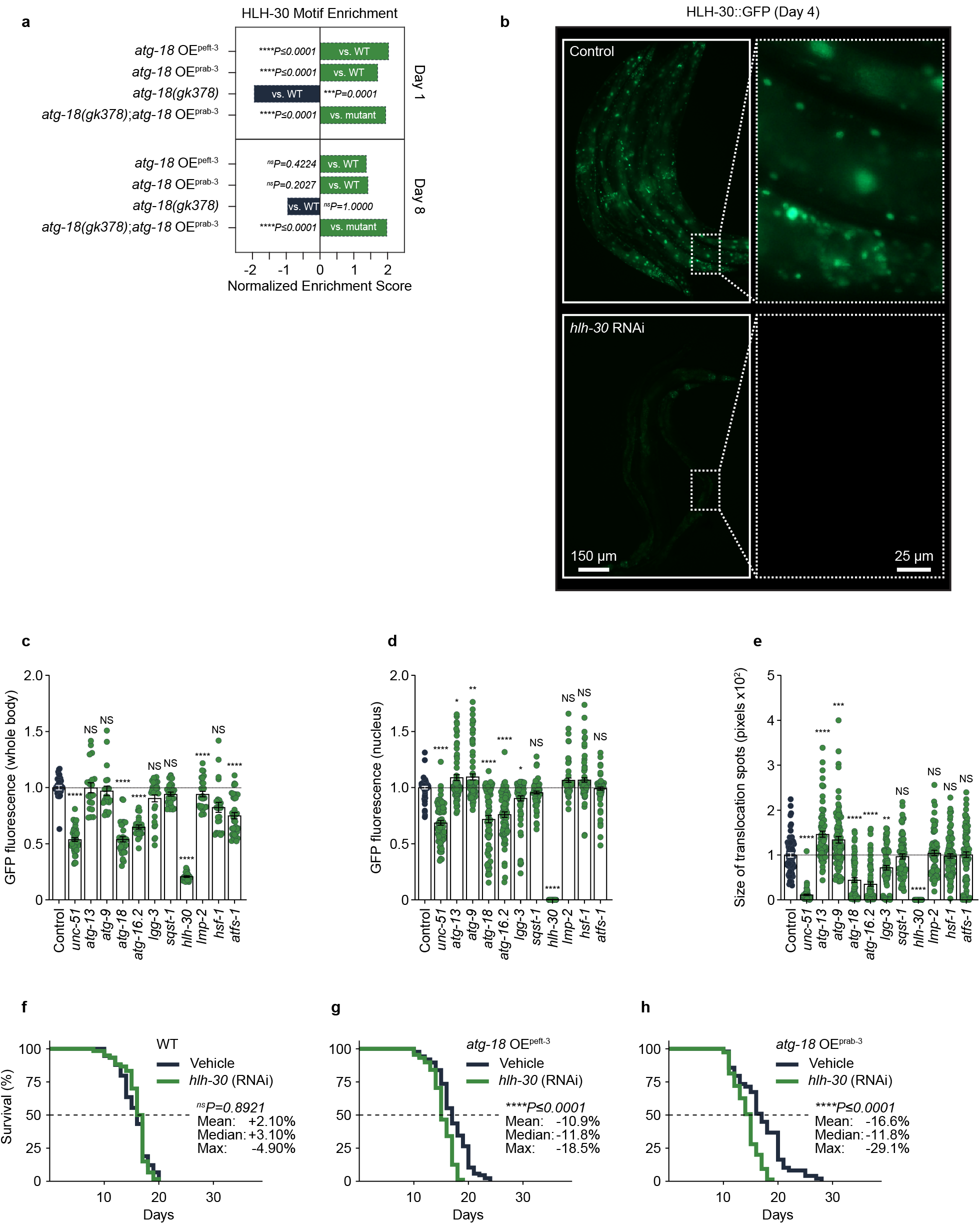
*hlh-30* knockdown blocks *atg-18* mediated lifespan extension and drives an age-dependant switch in *C. elegans*. **a**, Motif enrichment analysis of *hlh-30* for different levels of *atg-18* at day 1 and day 8 of adulthood. **b**, Representative images of HLH-30::GFP level with the RNAi knockdown of an array of genes involved in autophagy and other proteostatic systems at Day 4 of adulthood after exposure to 37°C for 1 h. **c**, Total fluorescent intensity per animal; each dot represents a single animal. **d**, Total nuclear fluorescent nuclear intensity; each dot represents a single animal. **e**, Size of translocation spots; each dot represents a single translocation spot. **f-h**, The impact of *hlh-30* knockdown in WT **(i)**, *atg-18* OE^peft-3^ **(j)**, and *atg-18* OE^prab-3^**(k)**. Data represent three biological replicates. ∼100 worms were used for each group/biological repeat. All quantitative data were shown in median ± S.E.M. Log-rank test was used for the statistics of lifespan data.

To experimentally address this potential link, we used an HLH-30::GFP reporter strain and performed RNAi knockdown of an array of several genes involved in autophagy and other proteostatic systems (**Fig. 3b-e**). Animals were exposed to 37°C for 1 h to induce nuclear translocation of HLH-30/TFEB. First, we found that RNAi knockdown of *atg-18, atg-16*.*2, lgg-3 and unc-51* abolished almost completely the nuclear translocation of HLH-30 (**Fig. 3e**) and also reduced the total levels of HLH-30/TFEB (**Fig. 3f-h**). It was previously shown that these four proteins can form a complex and interact with each other^50^. Hence, these data might suggest the potential regulation of HLH-30/TFEB levels by the autophagic protein complex independently of autophagy in *C. elegans*. To address whether interaction between *hlh-30* and *atg-18* is essential for the *atg-18* induced lifespan extension, we knocked down *hlh-30* through RNAi in the *atg-18* OE^peft-3^ and *atg-18* OE^prab-3^ nematodes, and observed a significant and similar reduction (-11.8% for both) of median lifespan in both strains, while *hlh-30* knockdown did not affect the lifespan in WT animals (**Fig. 3f-h**). Thus, our data indicate that *atg-18/WIPI2* regulates lysosomal biogenesis and motility with its lifespan extension capacity dependent on HLH-30/TFEB.

### ATG-18 impacts lysosomal number

Given that HLH-30’s mammalian orthologue, TFEB, is a master regulator of lysosomal biogenesis and autophagy, and that lysosome number correlates with age^51^, we asked whether *atg-18* influences lysosomal function. To address this question, we knocked down *atg-18* by RNAi in the NUC-1::CHERRY reporter (endonuclease that locates within lysosomes) to quantify lysosomal number at different ages^51^. Although lysosome number was unaffected at D1, *atg-18* knockdown led to a clear reduction in lysosome number by D4 compared to control animals (**Fig. 4a,b**). To further validate these findings, we performed LysoTracker Red staining, which labels acidic cellular compartments, including lysosomes, in strains the overexpressing *atg-18*. We found that both *atg-18* OE strains exhibited increased LysoTracker signal intensity; however, only the *atg-18* OE^prab-3^ animals reached statistical significance (**Fig. 4c**). Consistent with our earlier findings, the *atg-18* mutant, with or without neuronal rescue, showed a trend toward reduced LysoTracker signal that did not reach significance. Given the observed variability, additional biological replicates will be required to conclusively assess these effects.

**Fig. 4.**
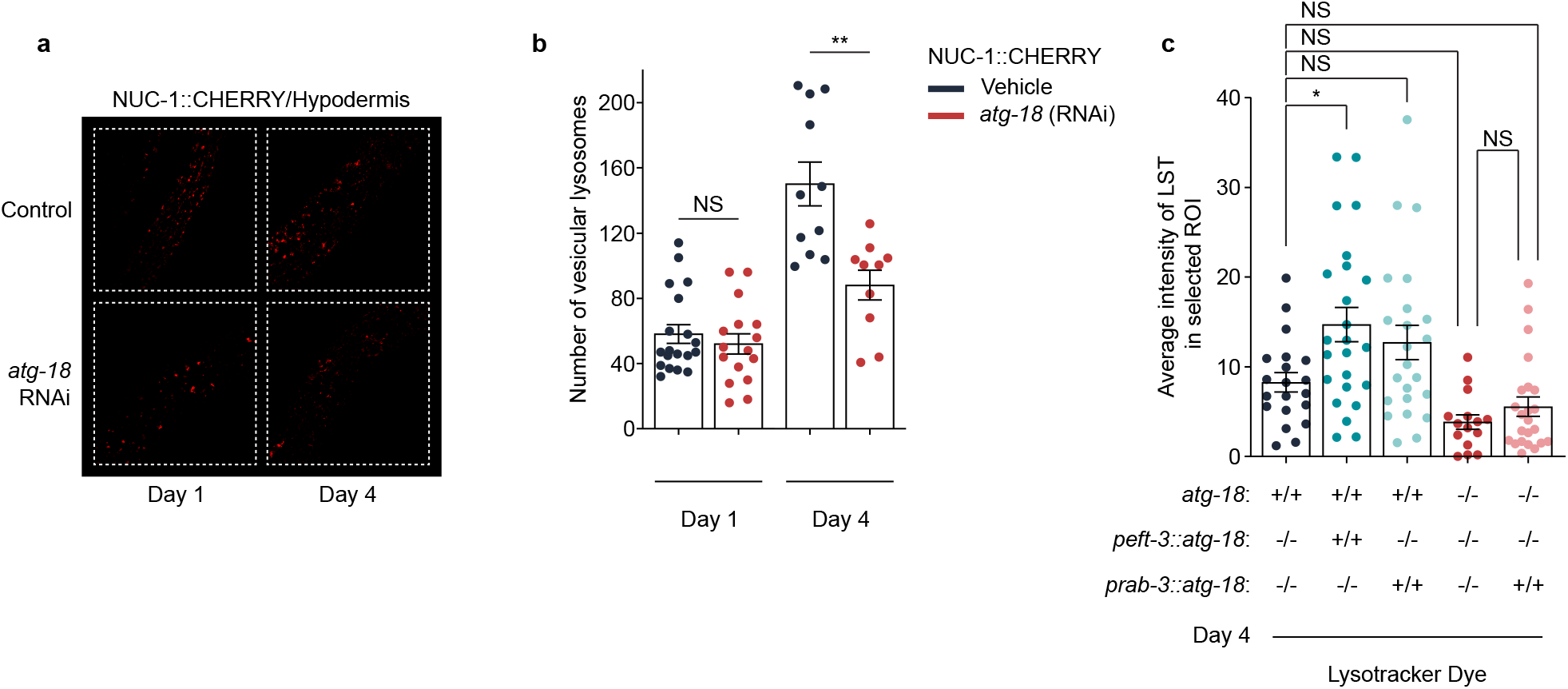
*atg-18* knockdown affects lysosomal number in *C. elegans*. **a**, Confocal fluorescence images of NUC-1::CHERRY reporter representing number of vesicular lysosomes with RNAi knockdown of *atg-18* at day 1 and 4 and (**b**) quantification of the number of vesicular lysosomes per animal. Each dot represents data pooled from three biological replicates. At least 10 animals were scored at each age. **c**, Quantification of Lysotracker Red staining. The number of lysosomes was quantified in the tail region of *C. elegans*.

## Discussion

In this study, we have further validate *atg-18* as a gene involved in health and longevity. In autophagy, two essential structural components, the autophagosome and the lysosome, fuse to form the autolysosome. While ATG-18 is a critical component in the biogenesis of autophagosomes, HLH-30 is a master regulator of lysosomal biogenesis. Here, we show for the first time a co-regulation between ATG-18 and HLH-30 on HLH-30 nuclear translocation, lysosomal number and lifespan extension. Moreover, ATG-18 ameliorates age-dependent transcriptional changes, overall suggesting the existence of a previously unrecognized ATG-18—HLH-30 autophagy—lysosome pathway that influences lifespan and healthspan.

Based on our results, it seems evident that *atg-18* acts in an antagonistic pleiotropic fashion under specific conditions (e.g. extreme temperature), where its benefits are accompanied by certain negative effects. We found that under extreme temperature conditions (4°C and 37°C) in young adulthood (D1), long-lived *atg-18* OE^peft-3^ and *atg-18* OE^prab-3^ worms died easily, while the short-lived *atg-18(gk378)* mutants were highly resistant (**Fig. 1**). Considering the importance of autophagy in heat shock resilience^52^, our findings may seem paradoxical; however, similar observations have been reported regarding HSF-1, the master regulator of the heat shock response^53^. Others have also shown that tissues/cells, like neurons, are particularly weak against heat stress early in life due to a delayed heat shock response^54^. These works may offer an explanation to our observations, where the lack of *atg-18* may result in the compensatory response of several pathways relevant for heatshock resilience; therefore be more prepared to face heat shock since the compensatory mechanisms are already activated. Altogether, similarly to HSF-1^53^, the role of ATG-18 in heat shock response may differ depending on the age of the organism: *atg-18* deletion animals were thermo-resistant in D1 but thermo-sensitive in D8 (**Fig. 1**).

We also showed a regulation of HLH-30 by ATG-18. RNAseq of strains with different *atg-18* expression revealed significant changes in HLH-30 motif enrichment (DNA sequence motifs that are enriched at sites bound by HLH-30), which is suggestive of the activity of this transcriptional factor. While HLH-30 is understood as a transcription factor that regulates a large array of autophagy-related genes, including *atg-18*^30^, we find that the levels of *atg-18* directionally regulate HLH-30 translocation and likely its activity. This previously unrecognized feedback mechanism may serve as a bridge between autophagosome formation and lysosomal biogenesis. Our findings align as well with early yeast studies showing that ATG-18 localization in proximity to acidic vacuoles in a pattern distinct from other autophagy proteins proteins^19^, as well as with more recent work in *C. elegans* indicating that ATG-18 exhibits pleiotropic functions that extend beyond canonical autophagy^36^. In this line, we also observe a bidirectional relationship between *atg-18* levels and lysosomal number, further supporting that HLH-30 is influenced by *atg-18*.

Furthermore, we have experimentally shown *in vivo* that the lifespan extension observed through *atg-18* requires the presence of *hlh-30*. Others have also shown that HLH-30 is essential for the lifespan extension of long-lived mutants such as *daf-2, eat-2* and *glp-1*^30^. Similarly, recent reports suggest that knockdown of *atg-18* seems to have a near identical effect on these long-lived strains^36^. We believe that our mechanistic findings complement and harmonize these findings, where ATG-18 and HLH-30 seem to be part of a key longevity pathway. Further studies could focus on how ATG-18 might regulate HLH-30 nuclear translocation. In normal condition, TFEB is phosphorylated (Ser142 and Ser211) by mTORC1 and ‘trapped’ by proteins 14-3-3; in conditions like starvation, the phosphatase calcineurin de-phosphorylates TFEB enabling its release from 14-3-3 and subsequent nuclear transport^23,24^. It is reported that mTORC1 regulates WIPI2 degradation via phosphorylating WIPI2 at Ser395 and directing its interaction with the E3 ubiquitin ligase HUWE1 for UPS-based degradation^55^. We thus speculate that WIPI2 may either weaken mTORC1’s phosphorylation of TFEB via competing with TFEB to bind mTORC1, and/or increase de-phosphorylation of TFEB by interacting with calcineurin or via other mechanisms. Further experiments are needed to understand the mechanisms.

Further work addressing the differential levels of *atg-18* across these overexpression strains, their spatial expression patterns, and how these differ from WT expression will help in further understanding the role *of atg-18*. In addition, it remains to be determined whether these effects are conserved in mammalian cells.

## Acknowledgement

We thank Professor Xiaochen Wang from fascilitating the NUC-1 reporter (XW5399). We thank Malene Hansen, Björn Schumacher, and their respective teams for their support and contributions to this project. The Ph.D. programmes of T.S.M. and A.A. were supported by HELSE SØR-ØST (#2021021) and the Civitan Norges Forskningsfond for Alzheimers sykdom (#281931), respectively. T.S.M. was also supported by Wellcome Leap’s Dynamic Resilience Program (jointly funded by Temasek Trust) (#104617) and the Cure Alzheimer’s Fund. E.F.F. was supported by Cure Alzheimer’s Fund (#282952),HELSE SØR-ØST (#2020001, #2021021, #2023093), the Research Council of Norway (#262175, #334361), Molecule AG/VITADAO (#282942), NordForsk Foundation (#119986), the National Natural Science Foundation of China (#81971327), Akershus University Hospital (#269901, #261973, #262960), the Czech Republic-Norway KAPPA programme (with Martin Vyhnálek, #TO01000215), the Rosa sløyfe/Norwegian Cancer Society & Norwegian Breast Cancer Society (#207819), HORIZON-TMA-MSCA-DN (#101073251, with Riekelt Houtkooper), and Wellcome Leap’s Dynamic Resilience Program (jointly funded by Temasek Trust) (#104617). R.X.A. is supported by Wellcome Leap’s Dynamic Resilience Program (jointly funded by Temasek Trust). JL was supported by grants from the Biotechnology and Biological Sciences Research Council (BB/W014890/1 and BB/T013273/1) and ST was supported by a PhD studentship from the Dunhill Medical Trust. G.L. was supported by research grants from Guangzhou Basic and Applied Basic Research Foundation (# 2023A04J1951), Guangdong Basic and Applied Basic Research Foundation (# 2022A1515111047), Fundamental Research Funds for the Central Universities, Sun Yat-sen University (# 24pnpy249). We are grateful to funding from MSD, the UK Dementia Research Institute (funded by the MRC, Alzheimer’s Research UK, and the Alzheimer’s Society), and Wellcome Leap’s Dynamic Resilience Program (jointly funded by Temasek Trust) to DCR. We are grateful for funding FDCT0052/2025/AJJ to H.M.S. and E.F.F.

## Methods

### *C. elegans* maintenance

Unless specified, the roundworm *C. elegans* animals were maintained at a temperature of 20 °C in NGM-containing plates. The nutrition of the animals consisted of the *Escherichia coli* strain OP50. The NGM was poured into empty plates, let dry for 2 days. At the second day of drying OP50 cultivated overnight at 37 °C was poured into the plates. The plates were then left to dry at room temperature once again for 2 days and from there were either used or stored at 4 °C for a maximum of a week for later use.

### Lifespan and pumping rate

Unless specified, lifespan assays were performed at 20 °C. At the first day of adulthood (the day after L4 is considered as adult D1 as we normally do)^56^, 30-35 worms were transferred into medium-size NGM plates (60 mm diameter) with 200 μL of *E. coli* (OP50) to obtain a synchronized adult population of around 100 animals. To avoid egg laying, the NGM also had a concentration of 75 μM of FUDR. Worms were examined every day and scored as dead if response to a touch was not recognized by the performer. For pumping assay, the number of pharyngeal muscles contractions in the synchronized worms were manually measured for 30 seconds at designated ages. Unless specified elsewhere, 3 biological repeats were performed.

### Assays on survival under different stress conditions: heat shock, cold shock, and starvation

Following the standard maintenance and lifespan assays, worms were heat shocked at 37 °C for 3 hours (with food) at designated ages. The animals were then returned to the maintenance temperature of 20 °C and scored as dead or alive 24 hours after they were removed from the hot environment. For cold shock assay, worms were at designated ages cold shocked at 4 °C for a period of 24 hours (with food). The animals were then returned to the maintenance temperature of 20 °C and scored as dead or alive 24 hours after they were removed from the cold environment. Regarding starvation assay, for L1 arrest starvation, worms were bleached with bleaching solution for synchronization. After washing, the eggs were placed into NGM plates without OP50. 5 days after, worms were scored either dead or alive^47^. For all three experiments: for each of them we had a total of 300 animals per experiment with 3 biological repeats.

### Pathology measurements

The experiment was performed per previous published method^42^. To assess intestinal atrophy, animals were cultured on NGM plates supplemented with 75 μM FUDR (5-Fluoro-2′-deoxyuridine, Sigma–Aldrich, F0503). Differential interference contrast (DIC) images were captured on Day 14 of adulthood using a Zeiss LSM 880 Confocal Microscope at 63x magnification. The severity of uterine tumors was evaluated by classifying the uterine mass either as separated eggs with distinguishable structures within or as a formed uterine mass where individual nuclei were no longer distinguishable. DIC images were taken on Day 1 and Day 4 of adulthood using Zeiss LSM 880 Confocal Microscope at 63x magnification. We had a total of 30 animals per experiment with 3 biological repeats.

### Fluorescence microscopy

To prepare worms for imaging analyses, HLH-30::GFP reporter strains were bleached, synchronized, and plated on NGM plates seeded with RNAi bacteria targeting the genes of interest from L1 stage, respectively. Worms at Day 1 and Day 8 of adulthood were anesthetized using 40 mM levamisole (Santa Cruz Biotech, cat. no. sc205730) and imaged at room temperature with a Nikon Eclipse Ti2 microscope at 10x magnification. For each condition, 30 worms were imaged, with each biological replicate containing at least 10 worms. GFP fluorescence was measured using ImageJ software (http://rsbweb.nih.gov/ij/) by analyzing the entire worm area. The same procedure was followed for the HLH-30::GFP reporter strain, but in this case, nuclear localization was quantified^30^. A distinct signal in the nucleus indicated nuclear localization, while the absence of a signal indicated non-localization.

### RNA-seq processing

Following our published protocol^57^, we collected adult D1 and D8 worms with 5 replicates/group for RNA-seq data generation. First we used Fastp v0.20.0^58^ with the following parameters to preprocess the raw fastq files: -g -x -q 30 -e 30 –trim_front 1. The preprocessed samples were mapped with Salmon v1.1^59^ and the parameters –validateMappings –gcBias – seqBias to a decoy-aware transcriptome reference of the WS281 transcriptome with a kmer size of 31. Tximport v1.14.2^60^ was used to combine the results of Salmon to the gene-level. We removed 4 outliers based on the expression values of *atg-18* for all analyses (FF36_d1_3, FF36_d8_3, ATG18_d1_4, ATG18_d8_3). Additionally we removed WT_d8_3 as this appeared to be a mislabelled d1 sample. edgeR v3.40.2^61^ in R v4.2.2 was used to compute the differentially expressed genes, including a correction for batch differences in the design matrix. The fgseaMultilevel function of fgsea v.1.24^62^ was used with nPerm=1000 to compute gene set enrichment analyses.

### Bit age predictions

Bit age was applied as previously described^48^. Briefly, to binarize the data, the median expression for each sample, after masking zeros genes, was calculated. Genes larger than the median were set to 1 and all other genes were set to 0.

### Transcription factor motif analysis

All CORE motifs for *C. elegans* from the 10th release (2024) of the JASPAR database were downloaded and used HOMER’s^63^ findMotifs.pl function for all motifs on all genes with a motif score of bigger than 7 to identify genes that are potentially regulated by the respective transcription factor. The list of genes were used as input for a GSEA analysis with fgsea v.1.24 as described above.

### Transcriptomic heatmap

The edgeR-normalized log-scale expression values of all 3026 genes that were at least significant (adjusted p-value <0.05) in 2 pairwise comparisons within the same timepoint. Plotted are the zscore normalized values of the median expression values within each sample and timepoint. The zscores are calculated across the 10 different conditions, i.e. a zscore of 0 if the mean expression value across all 10 conditions. The heatmap is plotted with Python’s Seaborn v0.11.0^65^ clustermap() function and the following parameters method=‘ward’, metric=‘correlation’.

### Statistics

For all experiments, unless otherwise specified, data were from 2-3 biological repeats. For most experiment, using Prism 10, the comparisons between two groups were performed using unpaired t test. When the assumption of normality was not met (determined by the Shapiro-Wilk test), we performed the Mann-Whitney U test. For multiple comparisons, we performed Ordinary one-way ANOVA. For egg-laying statistics Python’s statsmodels v0.11.1 mixedlm() function was used. We used a mixed-effects linear model to analyze the relationship between egg production, strain, and days of adulthood, while accounting for batch effects with the following setup: mixedlm (“eggs ∼ strain*day_adulthood, groups=‘Batch’, re_formula=∼’day_adulthood’). The model included fixed effects for strain, days of adulthood, and their interaction, and the random effect for batch with a random slope for days of adulthood, i.e. the effect of days of adulthood on egg production is allowed to vary across different batches.

---

**Supp. Fig. 1.**
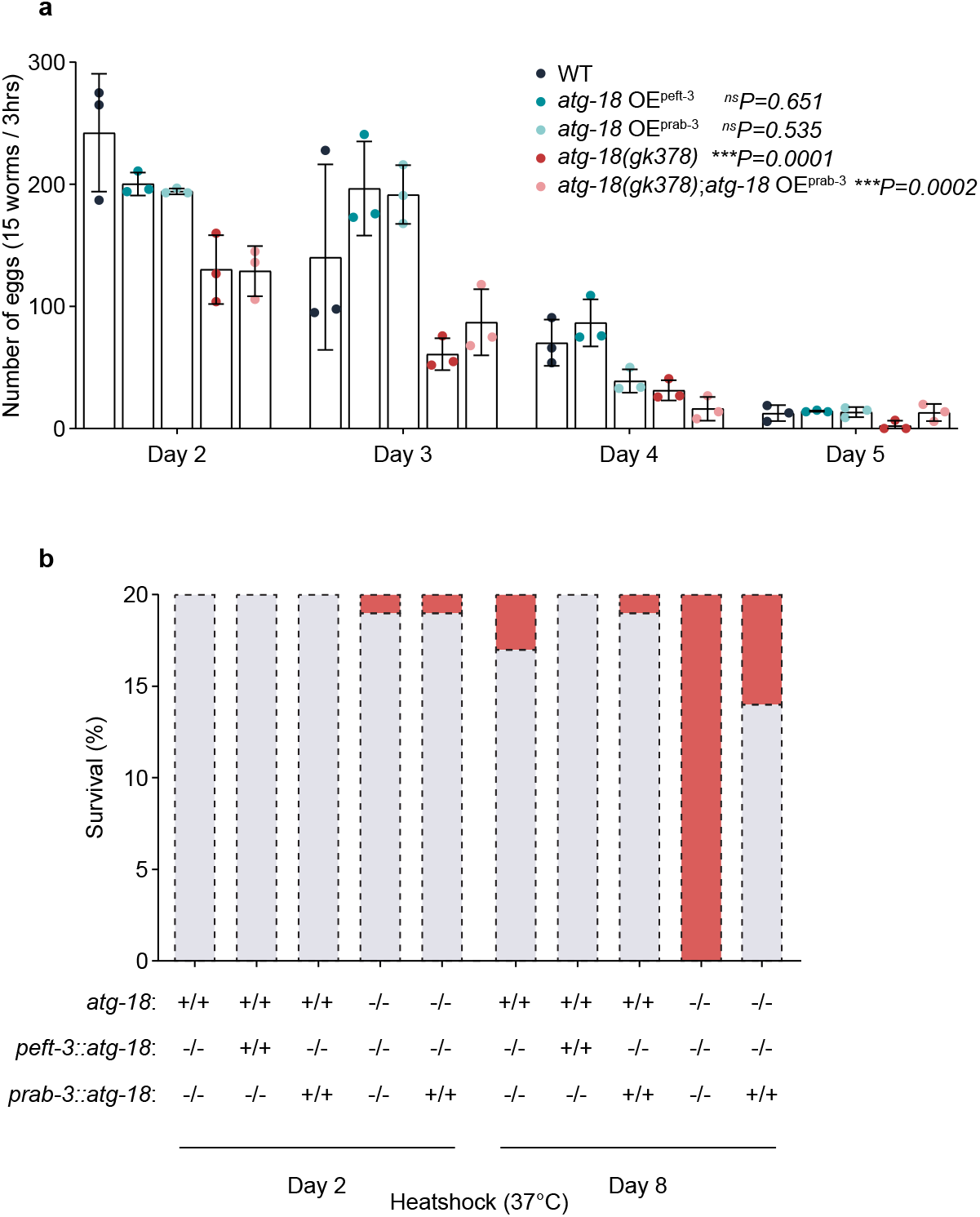
Additional healthspan and survival parameters of *C. elegans* under with different levels of *atg-18*. **a**, Fecundity changes at days 2, 3, 4 and 5 of adulthood. Each dot represents one biological repeat. At least 10 animals were scored in each strain at each age. **b**, Number of uterine tumors for different atg-18 level. Each graph represents data pooled from three biological replicates with at least 10 animals scored in each strain at each age. The data show quantification for 20 selected animals in each strain.

**Supp. Fig. 2.**
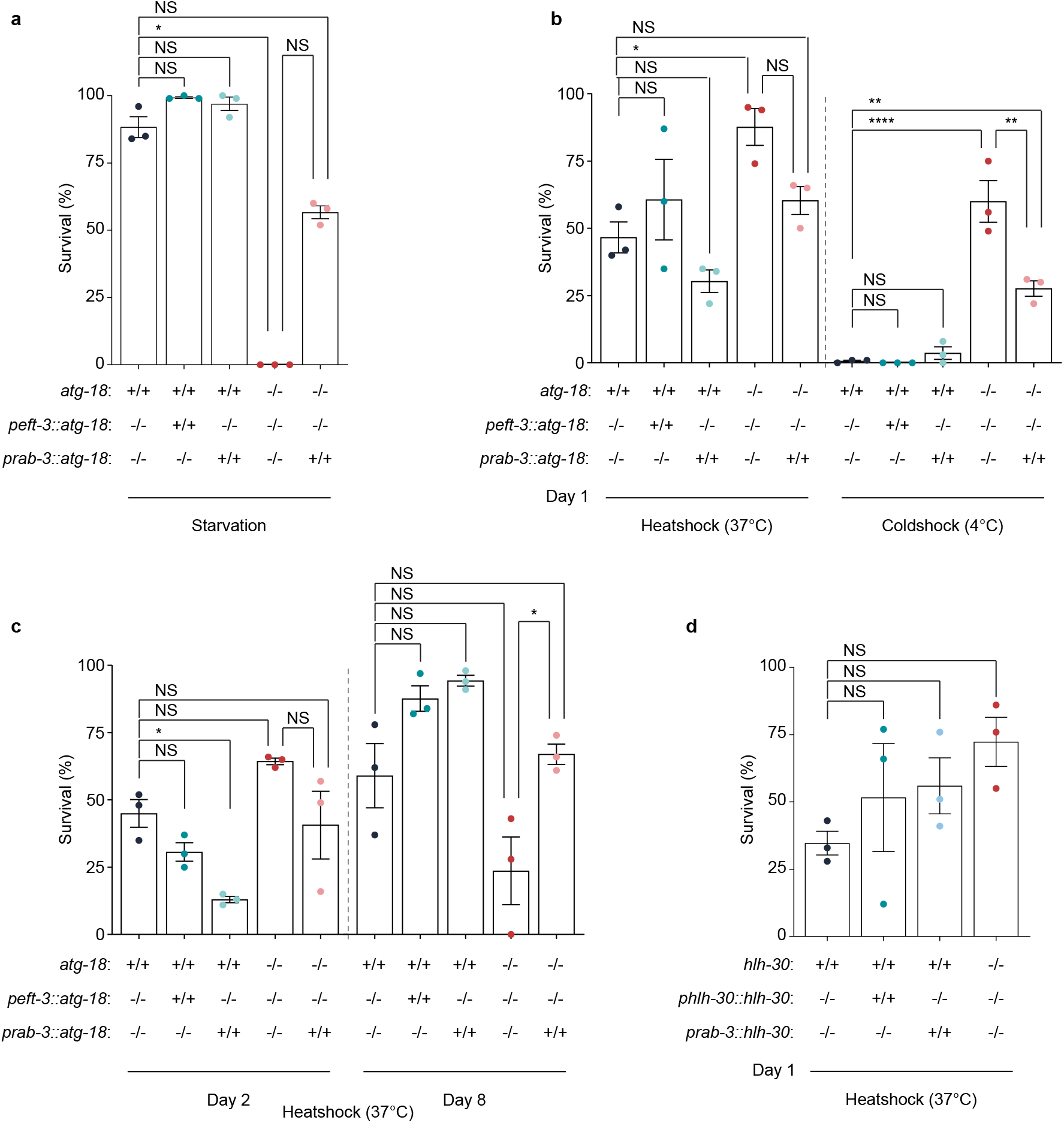
Stress and survival assays of *C. elegans* under with different levels of *atg-18*. **a-d**, Survival of *C. elegans* after 7 days of starvation at L1 arrest stage **(a)**, after a 24-hours cold shock at 4°C at day 1 of adulthood or after a 3-hour heat shock at 37°C at day 1 **(b)**, at day 2 **(c)** and day 8 **(d)**. Each dot represents a single biological repeat, with each biological repeat consisting of around 100 animals.

